# Membrane Proteins Significantly Restrict Exosome Mobility

**DOI:** 10.1101/196691

**Authors:** Mikhail Skliar, Vasiliy S. Chernyshev, David M. Belnap, Samer M. Al-Hakami, Inge J. Stijleman, Rakesh Rachamadugu, Philip S. Bernard

**Affiliations:** Department of Chemical Engineering, University of Utah; Nano Institute of Utah; Skolkovo Institute of Science and Technology, Moscow, Russia; Life Sciences Center, Moscow Institute of Physics and Technology, Moscow, Russia; Departments of Biology and Biochemistry, University of Utah; Huntsman Cancer Institute, University of Utah; Department of Pathology, University of Utah

**Keywords:** Exosomes, Hydrodynamic mobility, Surface proteins

## Abstract

Exosomes are membrane nanovesicles that intermediate cell-to-cell signaling through the transfer of their molecular cargo. The exosomes’ small size facilitates rapid migration through the extracellular matrix and into and out of circulation. Here we report that the mobility of the exosomes is much lower than would be expected from the size of their membrane vesicles. The difference is broadly distributed and caused by surface proteins, which significantly impede exosome migration. The observed wide range in the mobility implies that a subpopulation of hydrodynamically small exosomes is more likely to participate in signaling. The extracellular environment amplifies the size-dependent hindrance to the exosomes migration. The significant contribution of surface proteins to the transport resistance make the exosome mobility a dynamic property that changes with the extracellular environment which affects the membrane protein conformation, glycosylation, specific, and non-specific surface adsorption.

## INTRODUCTION

Exosomes are actively secreted by cells and can be isolated from all biological fluids, including blood, urine, and saliva. Exosomes are distinguished from other extracellular vesicles (EV), such as ectosomes,^1,2^ by biomarkers of the late endocytic pathway^3–5^ and by small size, typically reported between 30–150 nm. These properties are often used to confirm that EVs obtained from biofluids or cell culture are exosomal in origin. Several exosome isolation techniques, such as size exclusion chromatography^6–8^ and ultrafiltration,^9,10^ are entirely based on exosomes’ small size.

The luminal and surface composition of EVs are derived from the cells that shed them.^11,12^ Exosomes’ membrane and surface cargo – such as membrane proteins, saccharide groups, and other membrane-bound and adsorbed molecules – preserve the same topological orientation as in plasma membrane.^4,13^ By fusing with proximal and distal cells, exosomes mediate intercellular signaling in health and disease. The biologically active molecules transferred by the exosomes from the parent to the recipient cells include surface and luminal proteins,^14,15^ membrane-bound microRNAs,^3,16,17^ and other compounds.^18–20^ A growing number of studies have associated exosomal signaling with tumor metastasis,^21–24^ cancer drug resistance,^25^ and the modulation of the immune response.^26–29^

For signaling to occur, it is necessary, but not sufficient, for a secreted EV to traverse the distance from the parent to a recipient cell. For paracrine signaling, the transport barrier to exosomal communication may be overcome by passive diffusion through extracellular space, while to reach distal targets an EV may need to enter and exit a biofluid circulation. For example, the convective transport of EVs by a circulating biofluid is likely required for the formation of organotropic pre-metastatic niches and metastasis to distal sites, as in the case of lung cancer exosomes internalized by bone cells.^21^

The smallest in size among all EVs, the exosomes have the highest mobility through the extracellular matrix and in circulating biofluids, which favors exosome-mediated paracrine and distant signaling.^30^ The mobility of the exosomes, characterized by their mean squared displacement with time, is proportional to their (self-) diffusivity. The hydrodynamic diameter of the exosomes is the measure of the resistance to their migration and is inversely proportional to their diffusivity. Only in a limited case of a spherical, smooth, and hard nanoparticle (NP) with the electrically neutral surface will the hydrodynamic diameter equal the geometric diameter of the sphere. For exosomes, which are elastic lipid bilayer particles^31^ with membrane-conjugated macromolecules^32^ and a negative zeta potential^33^, these conditions do not hold. Consequently, the resistance to the exosomes migration should be higher than implied by the size of their membrane vesicle.

The transport barriers to exosome dissemination are poorly understood. Here, we examine the impact of surface proteins and macromolecules anchored by them, on the mobility of the exosomes. To this end, the hydrodynamic diameter of exosomes, isolated by precipitation from the growth medium of MCF7 (ER+) breast cancer cells, was analyzed before and after enzymatic digestion of the surface proteins and compared with the estimated size of the membrane envelope. The thickness of the coronal layer, equal to the difference between hydrodynamic and vesicle diameters, was used to quantify the resistance to the exosome migration caused by surface-conjugated macromolecules. We found a substantially impeded migration of exosomes prior to digestion relative to the expectations based on the size of their vesicles. Furthermore, the hydrodynamic diameters were found to be broadly distributed relative to a narrower range of vesicle sizes, indicating a widely varying coronal thickness. The enzymatic cleavage of surface proteins increased the mobility and shifted it closer to the expected range for the size of membrane envelopes. The significant and heterogeneous impact of the coronal layer on the reduction in the exosome mobility was then confirmed by analyzing the hydrodynamic and the membrane diameters of exosomes isolated from serum of a pancreatic cancer patient and a healthy individual. Several implications of these findings are discussed. The exosomes with higher mobility are more likely to reach a cellular target for signaling to occur, while those with large hydrodynamic diameters are more likely to accumulate and degrade in the extracellular matrix. Our results indicate that the difference in mobility impacting the signaling outcome is strongly impacted by a widely-distributed thickness of the coronal layer. Furthermore, the significant contribution of surface proteins to the transport resistance and the influence of the extracellular environment on protein conformation, specific and non-specific adsorptions, proteolytic degradation, and protein glycosylation indicate that the hydrodynamic size of the exosomes should be viewed as a dynamic property that changes with time and migration.

## RESULTS

### Characterization of isolated EVs

The size, shape, membrane morphology, and biomarker expression on extracellular particles isolated from the MCF7 cell growth medium are summarized in Fig. 1 The hydrodynamic size distribution (Fig. 1a) has the shape, the mean (equal to 109± 6 nm), and the mode (82± 3 nm) diameters consistent with previous reports.^34,35^ A small population of larger particles in Fig. 1a, likely consisting of co-precipitated ectosomes, was excluded from subsequent analysis. Morphological characterization by cryogenic transmission electron microscopy (cryo-TEM) shows the expected membrane vesicles of globular shape (Fig. 1b). The total protein concentration in the MCF7 exosome isolate containing approximately 10^12^ particles/mL was 3.5 mg/mL (includes the contribution of exosomal and background proteins co-isolated from the growth medium), which is typical for precipitation-based exosome isolation,^36^ and too low for the background proteins to agglomerate.^37^ The expression of several exosomal surface biomarkers was characterized by a reverse dot blot antibody array. Fig. 1c shows positive staining for Annexin A5 (ANXA5), tumor susceptibility gene 101 (TSG101), and tetraspanins CD63 and CD81 (further details in Supporting Information, SI). The staining for cisGolgi matrix protein GM130 (characterizes contamination of the exosome preparation) was negative. The abundance of luminal onco-microRNA, miR-21, previously reported to be highly expressed in the MCF7 exosomes,^38^ was confirmed by digital droplet PCR (Fig. S1 in SI).

**Figure 1.**
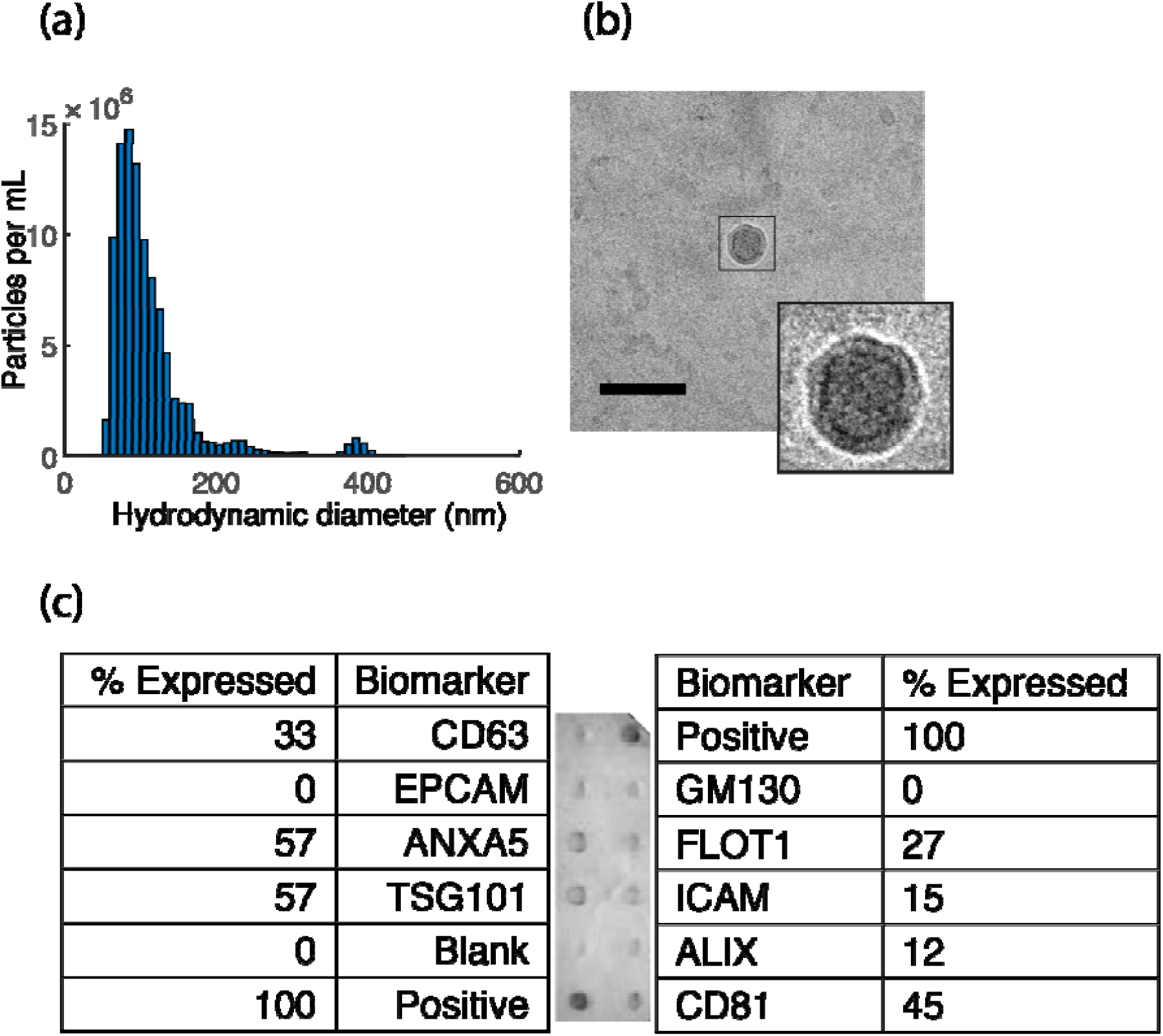
Size, morphology, and surface biomarkers of isolated MCF7 EVs. (a) Distribution of hydrodynamic diameters of isolated EVs. The group represented by the peak at 400 nm was excluded from further analysis. (b) CryoTEM image shows the expected globular morphology and the vesicle membrane (inset). The scale bar is 100 nm. (c) The quantification of the antibody array shows positive staining for several exosome biomarkers.

### AFM measurements of vesicle size

The membrane envelope of the hydrated exosomes, electrostatically immobilized on mica surface, was characterized by atomic force microscopy (AFM). The representative tapping-mode height and phase images of the surface-bound exosomes maintained in PBS are shown in Fig. 2a and b. The section of the height image along the line crossing three different exosomes (Fig. 2c) and the height image of one of the exosomes in Fig. 2d show severe shape distortion from the expected globular geometry^39–42^ caused by the electrostatic attraction of the exosomes, known to have a negative zeta potential, to the mica surface modified to be positively charged. The highly oblate membrane geometry was analyzed individually for 561 hydrated exosomes. The peak height the exosomes protrude above the mica surface was, on average, 7.9±3.1 nm (refer to Fig. 2c for an illustration). The surface area occupied by an immobilized exosome was approximated by a circular area with the diameter equal to twice the mean distance from the particle’s "center of mass "to its boundary. On average, this diameter was equal to 69.6±19.7 nm. Both of these averages are in the range of values previously reported for exosomes of different origin.^43,44^ Fig. 3a summarizes the results of the analysis in the form of empirical probability density functions (pdf) of the peak particle elevations above the substrate and the mean diameters of the occupied surface area.

**Figure 2.**
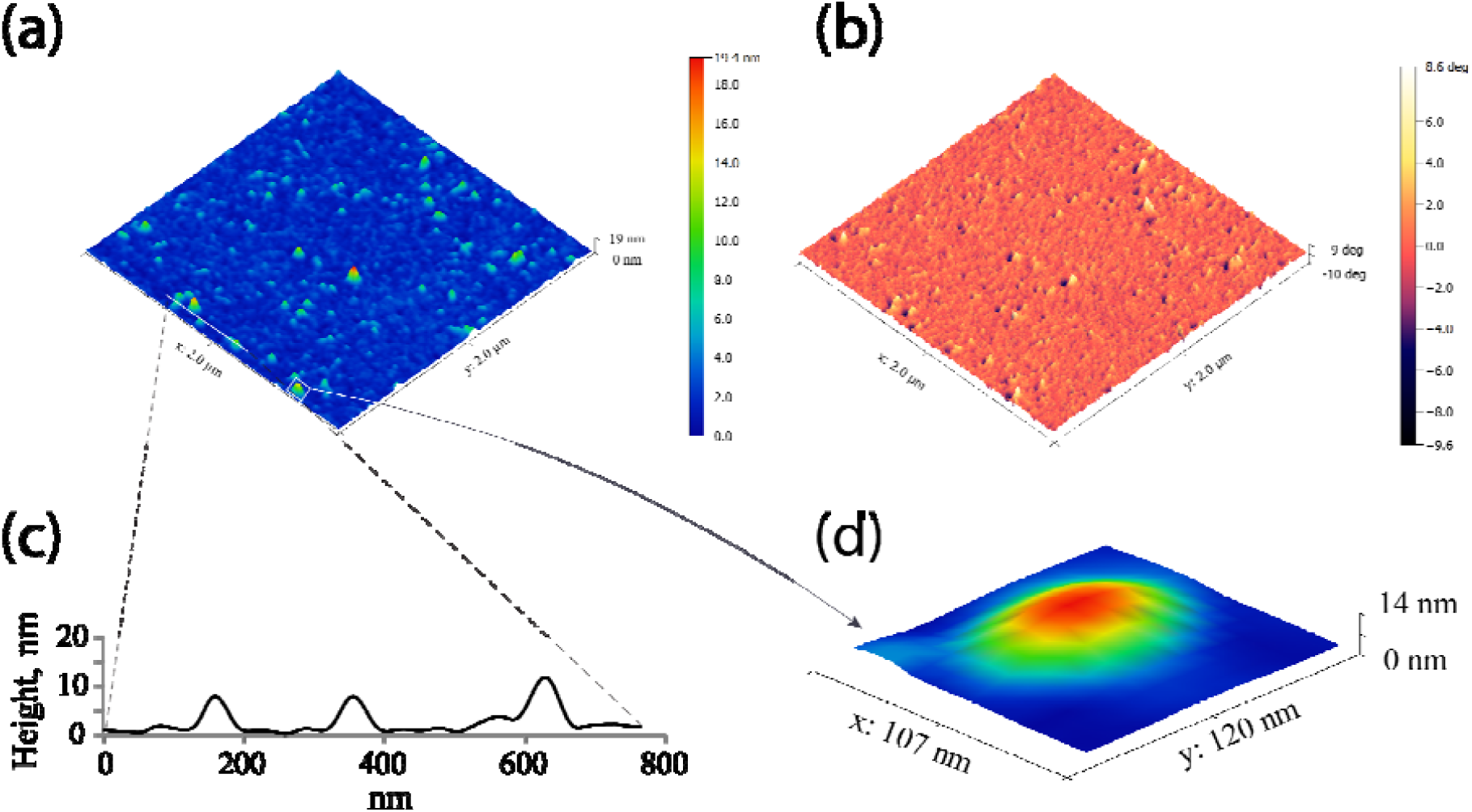
(a) Typical AFM height image of hydrated exosomes immobilized on the mica substrate. (b) The AFM phase image, sensitive to localized stiffness variations of soft samples, is consistent with the height image. (c) Detailed height data for several exosomes cross-sectioned by the line in panel (a). (d) A highly-distorted flattened shape of the immobilized exosome is caused by electrostatic interaction with a charged mica surface.

**Figure 3.**
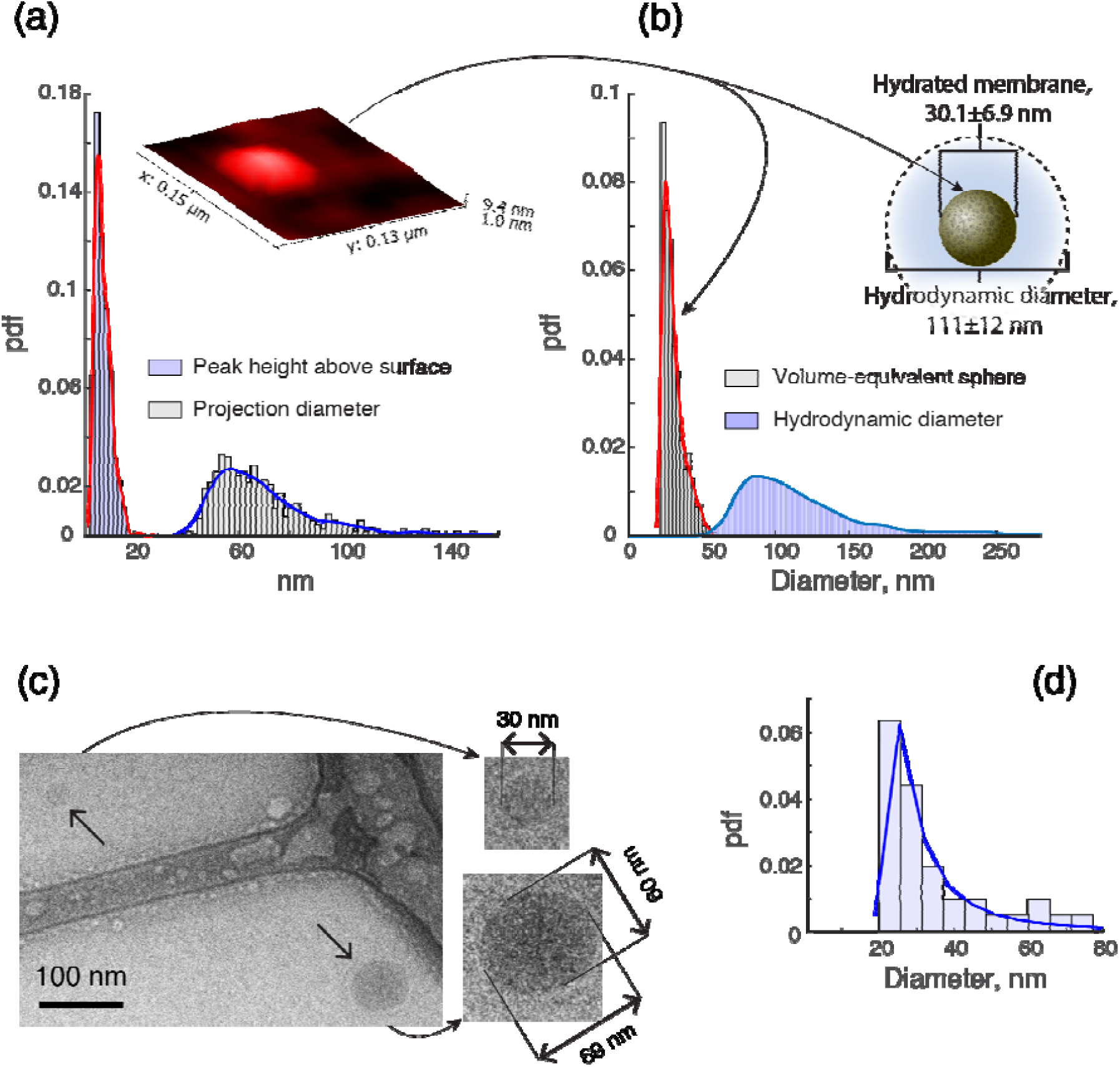
Exosome diameter and volume measurements. (a) AFM height measurements (inset) illustrate the highly oblate shape of exosomes after their electrostatic immobilization on the mica surface. The distribution of peak heights above the surface (red curve, blue fill) has an average equal to 7.9 nm. The area occupied by immobilized exosomes was characterized by the projection diameter, equal to twice the mean distance from the "center of mass" to the boundary of an exosome on the mica surface. The mean projection diameter for the obtained distribution (blue curve, gray fill) is 69.6 nm. (b) The size of the spherical membrane envelope for the exosomes in the solution was approximated to match the volume enclosed by the oblate shape of the surface-immobilized exosomes as measured by AFM. The distribution of the volume-equivalent spheres (red curve, gray fill) indicates a substantially smaller vesicle size than implied by the corresponding measurements of the hydrodynamic diameters by NTA (blue curve, blue fill). The thickness of the exosomal corona (inset) is estimated as the difference between hydrodynamic and membrane vesicle sizes. (c,d) Cryo-TEM imaging (c) confirms the globular shape of the hydrated exosomes in the solution and a small vesicular size (d) compared to their hydrodynamic mobility.

We used the AFM height data to calculate the volume enclosed by the membrane envelope of each surface-bound exosome. The obtained volume for each analyzed exosomes was mapped into the corresponding volume-equivalent sphere (insets of Fig. 3), the diameter of which provided size estimates of the membrane envelope in an innate globular shape. The obtained diameters of volume-equivalent spheres for the hydrated exosomes are characterized by the pdf shown in Fig. 3b (red trace). The ensemble average of this distribution is equal to 30.1±6.9 nm (Table 1), which is substantially smaller than the average of the hydrodynamic diameters (equal to 111±12 nm, Fig. 3b).

**Table 1.** AFM characterization of exosome sizes.

|  | *Analyzed particles* | *Average vesicle* |  | *Average diameter of volume-equivalent sphere* |
| --- | --- | --- | --- | --- |
|  |  | *Peak height above surface* | *Projection diameter* |  |
| *Hydrated sample* | 561 | 7.9±3.1 nm | 69.6±19.7 nm | 30.1±6.9 nm |
| *Desiccated sample* | 400 | 3.8±1.3 nm | 100.2±24.0 nm | 29.3±5.2 nm |

After allowing the immobilized exosomes to dry, the mica surface was rescanned by the AFM (Fig. S2). The desiccation further distorts the shape of the immobilized exosomes, making it even more oblate by reducing the height above the surface and expanding the occupied surface area (Table 1). Somewhat surprisingly, the volume enclosed by the desiccated vesicles remains almost the same as in the hydrated state.

### Cryo-TEM imaging and vesicle sizing

After rapid vitrification of unstained exosome sample, we used cryo-TEM to directly visualize and size the vesicles in the MCF7 exosome preparation. he observed globular appearance of the exosomes in the solution and the vesicle sizing procedure are illustrated in Fig. 3c. The average diameter of the imaged vesicles was 34.2 nm (Fig. 3d), which is consistent with the AFM-based estimate and an independent confirmation of small vesicular particles having a much larger hydrodynamic diameter. The slight difference between the cryo-TEM and AFM estimates may be explained by a higher mobility of smaller exosomes, which reach the mica surface more readily during the incubation period and skew the AFM results towards smaller sizes.^45^

### Impact of digestion on exosome mobility

We have previously hypothesized^39^ that membrane-conjugated macromolecules are responsible for the observed significant difference between the membrane and hydrodynamic sizes of the exosomes. We tested this hypothesis directly by measuring the hydrodynamic size of MCF7 exosomes before and after enzymatic proteolysis of their membrane proteins. The digestion protocols summarized in Table 2 utilized enzymes with different proteolytic specificity and varied the treatment duration.

**Table 2.** Digestion protocols.

|  | *Enzyme* | *Incubation* | *Quenching* |
| --- | --- | --- | --- |
| Protocol A | Trypsin | 20 min at 37°C | yes |
| Protocol B |  | 60 min at 37°C | no |
| Protocol C | Proteinase K | 60 min at 37°C | yes |
| Protocol D |  | 60 min at 37°C | no |

The size distributions of hydrodynamic diameters before and after digestive treatments were measured by the NTA. The results show an increase in exosome mobility by as much as 50% after enzymatic cleavage of their surface proteins (Fig. 4 and Table 3). We ruled out several factors potentially contributing to the observed reduction in hydrodynamic sizes. First, it was found that enzymatic treatments did not affect the size of the membrane envelopes, which remained within the same 20 to 40 nm range before and after the treatment (Fig. S3a-c). Second, the effect of solubilized proteins on the viscosity is negligible at the protein concentration in our sample.^46^ Consequently, the proteolysis of background proteins has no impact on the viscosity of the solution and the mobility of particles in it. Finally, a small reduction in the particle concentration after the digestion is insufficient to explain the observed large shift in hydrodynamic diameters towards smaller sizes even if in the worst-case scenario of only the largest particles lost from the ensemble. It was likely caused by nonspecific adsorption of exosomes to interfaces during sample transfer steps associated with the digestion and the NTA analysis before and after its completion.

**Table 3.**
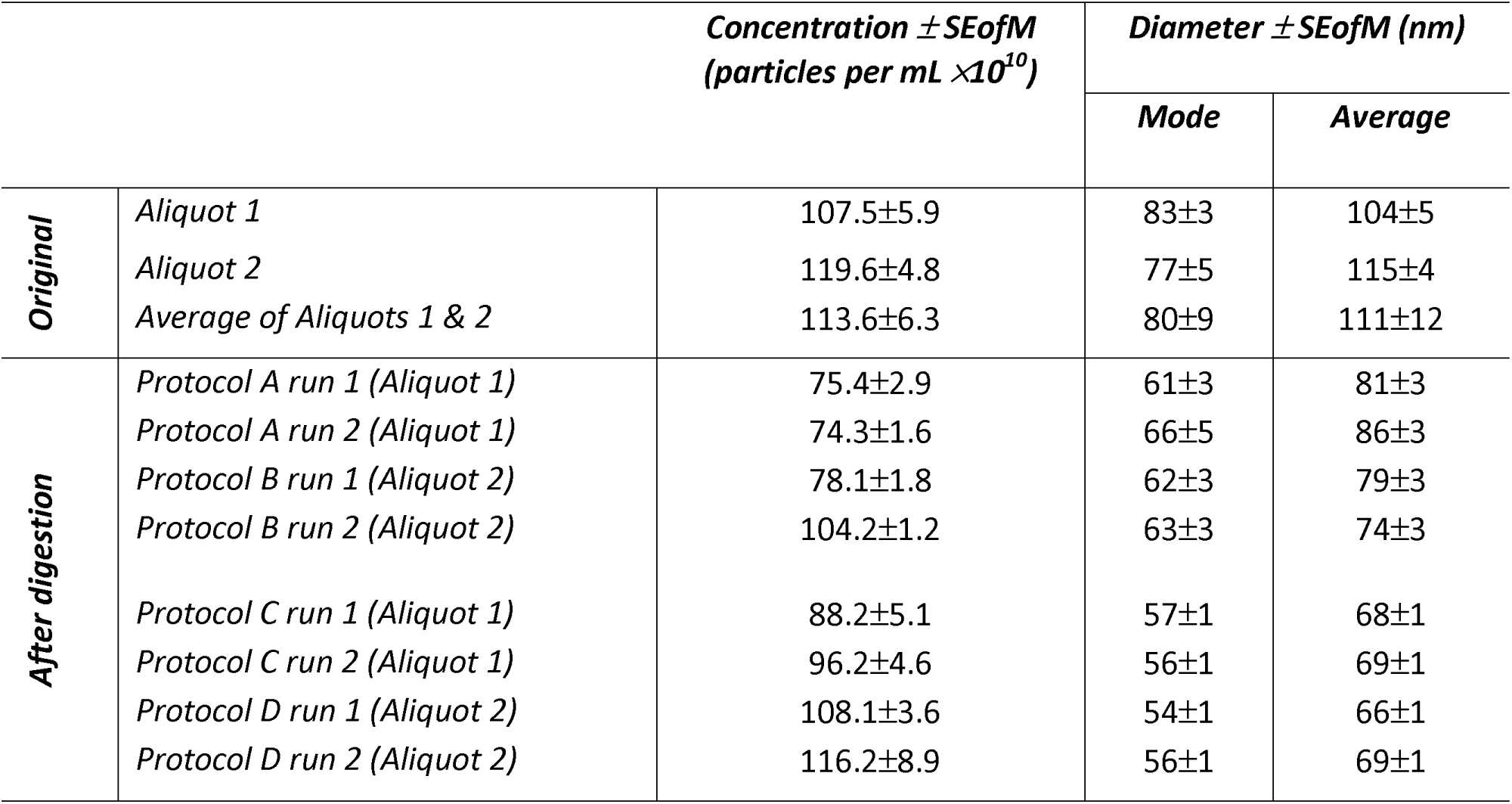
Concentration and hydrodynamic size (± standard error of the mean, SEofM) of exosomes before and after digestion.

**Figure 4.**
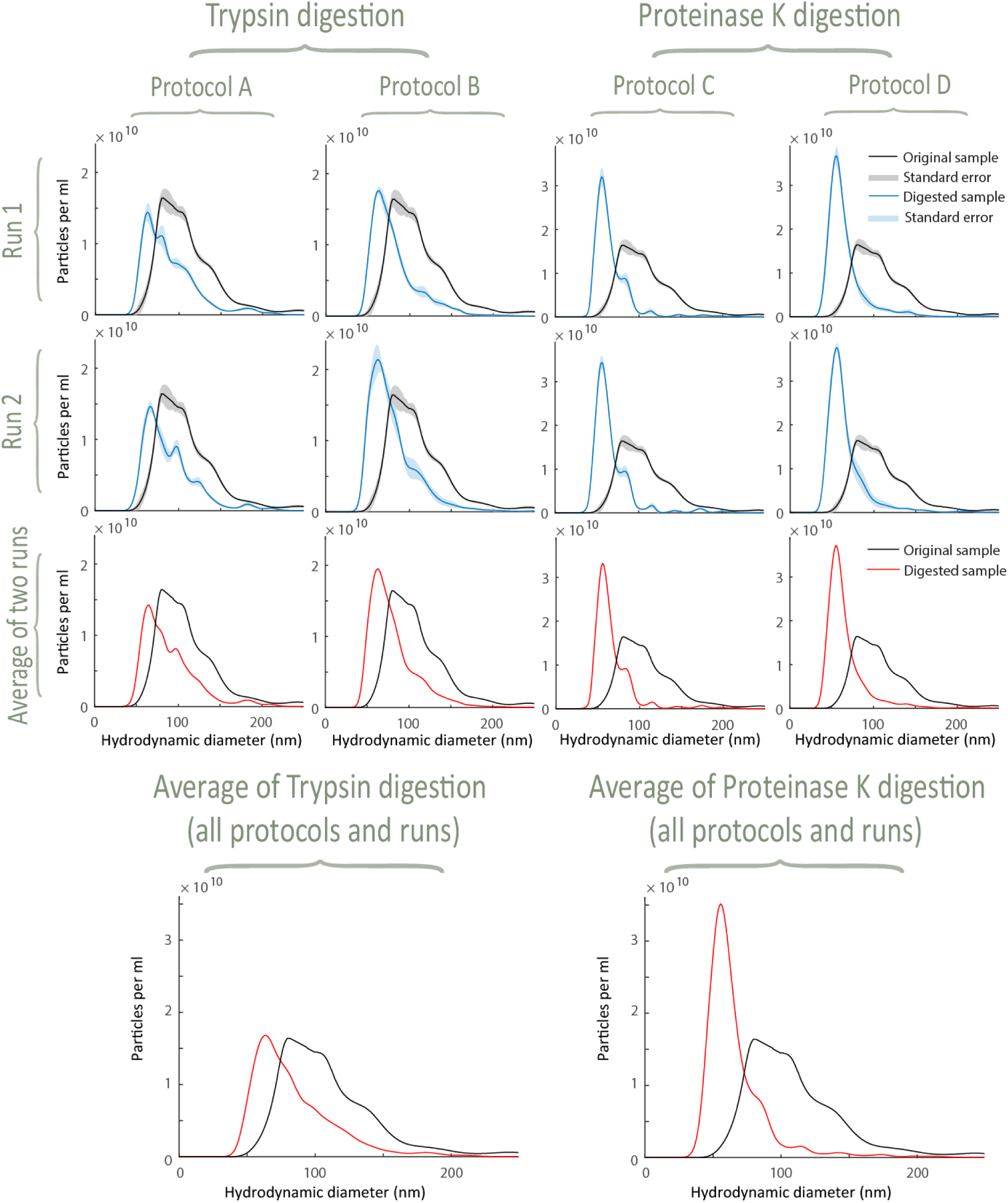
Hydrodynamic diameters are reduced substantially by enzymatic proteolysis of surface proteins. After the digestion, the sizes are shifted closer to the size of the membrane envelopes estimated from the AFM measurements and cryo-TEM imaging. The shift is especially pronounced after the less specific protease K digestion. After cleaving the surface proteins, the hydrodynamic size distribution is relatively narrow, which points to the heterogeneity in the macromolecular surface decoration as the source of widely different mobility of untreated exosomes.

Trypsin proteolysis preferentially cleaves the proteins at lysine and arginine locations and leaves intact segments of surface proteins void of these α-amino acids. After a relatively specific trypsin digestion, the mode diameter decreased to 64 nm (average of Protocols A and B in Fig. 4) from the original 80 nm. Proteinase K has a broader activity and cleaves peptide bonds of hydrophobic, aliphatic, and aromatic amino acids. As a result, the undigested fragments of the surface proteins are shorter than obtained after trypsinization, leading to the higher mobility of PK-treated exosomes. Quantitatively, the mode of hydrodynamic diameters after PK digestion following Protocols C and D reduced from 80 to 56 nm (Fig. 4).

The indiscriminate PK proteolysis shifts the hydrodynamic diameters of the exosomes into the range of sizes that partially overlaps with the size of the membrane envelopes characterized by cryo-TEM and AFM (Fig. 5). The remaining difference may be explained by the presence of short fragments of surface proteins that survived the digestion. The duration of enzymatic treatments in our experiments had little effect on the hydrodynamic sizes, likely because the incubation was sufficiently long in all protocols to achieve a near completion of the digestive process.

**Figure 5:**
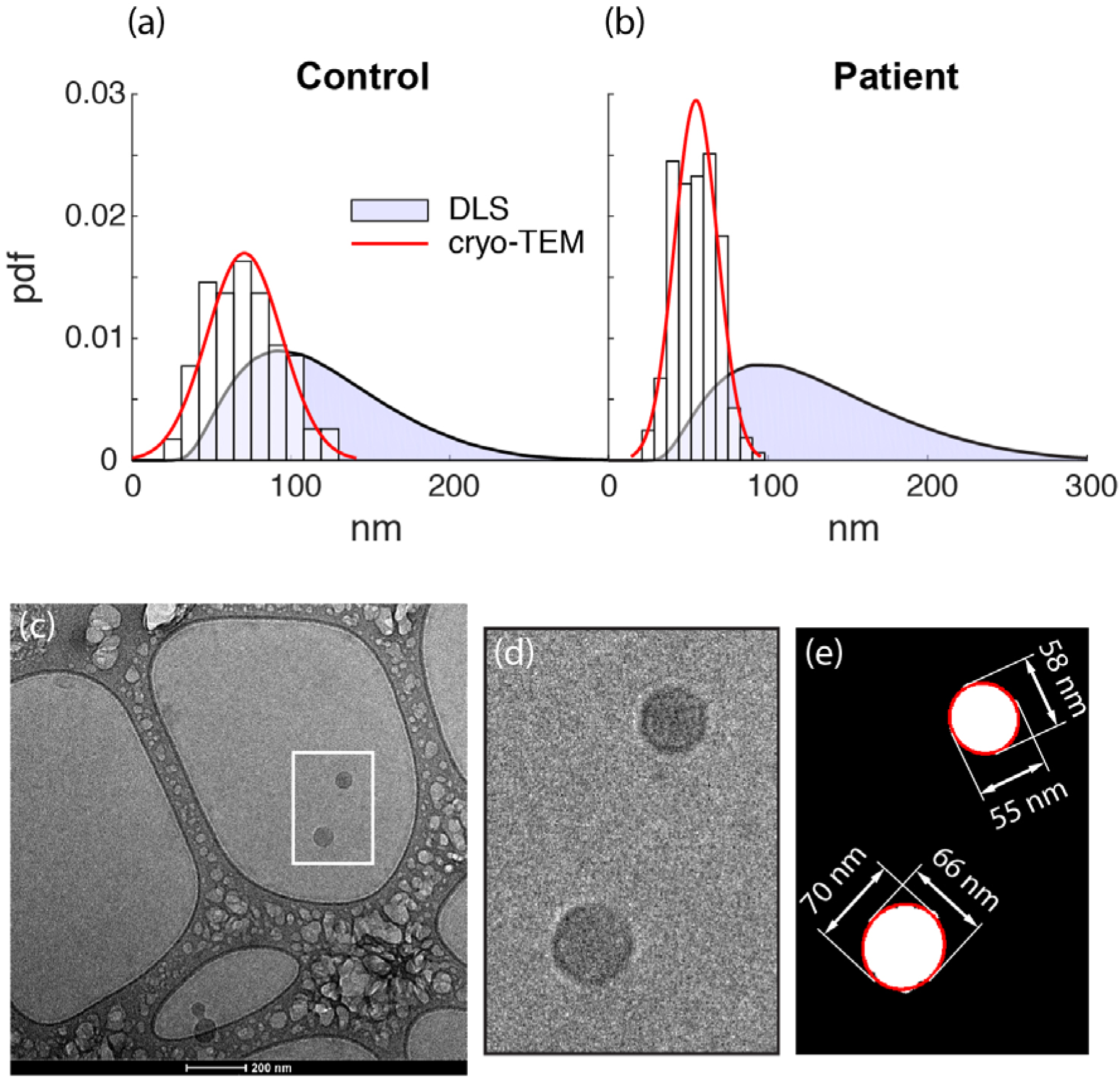
Sizes of human sera exosomes. (a,b) Hydrodynamic diameters of exosomes isolated from serum of a healthy control (a) and a pancreatic cancer patient (b) were obtained by dynamic light scattering. Measurements of vesicle sizes obtained from cryo-TEM images are shown for comparison. A representative cryo-TEM image (c) of the exosomes isolated from serum of a healthy individual illustrates their globular shape. (d) Extract of the micrograph shown in (c). (e) Computer analysis of cryo-TEM images was used to size the vesicles. Images (d-e) demonstrate the functionality of the image analysis software used to size the vesicles by calculating the geometric mean of the major and minor lengths of the ellipses fitted to each vesicle.

The hydrodynamic sizes after the digestion (Fig. 4) are distributed more uniformly. The distribution is particularly narrow after the less specific proteinase K treatment, which closely "shaves" the membrane envelope, leaving behind only short remnants of surface proteins. We, therefore, conclude that the broadness in the distribution of the hydrodynamic sizes is the consequence of the heterogeneity in the macromolecular surface decoration of the narrowly distributed exosomal vesicles.

### Vesicle vs. hydrodynamic size of serum exosomes

The validity of the described observations beyond MCF7 exosomes was examined next. To that end, the exosomes isolated from the serum of a pancreatic cancer patient and a healthy control were sized.^39^ Computer analysis of cryoTEM images (Fig. 6c-e) was used to size the vesicles in sera samples. The vesicle size distributions in Figs. 6a-b give the pdf of the geometric mean of the major and minor lengths of the ellipses fitted to each identified vesicle. The average vesicle diameters are 71±24 and 55±14 nm for the control and patient samples, respectively. The hydrodynamic diameters for both sera samples, measured by dynamic light scattering (DLS), are distributed broadly (Figs. 6a,b). The mode and mean for the control were found to 91 and 119±47 nm, respectively, whereas for the patient sample they were 92 and 130±55 nm. Therefore, similarly to the MCF7 cell culture case, the hydrodynamic diameters of serum exosomes are much larger than their vesicle diameters, which indicates the presence of a large macromolecular corona responsible for the reduced mobility.

**Figure 6:**
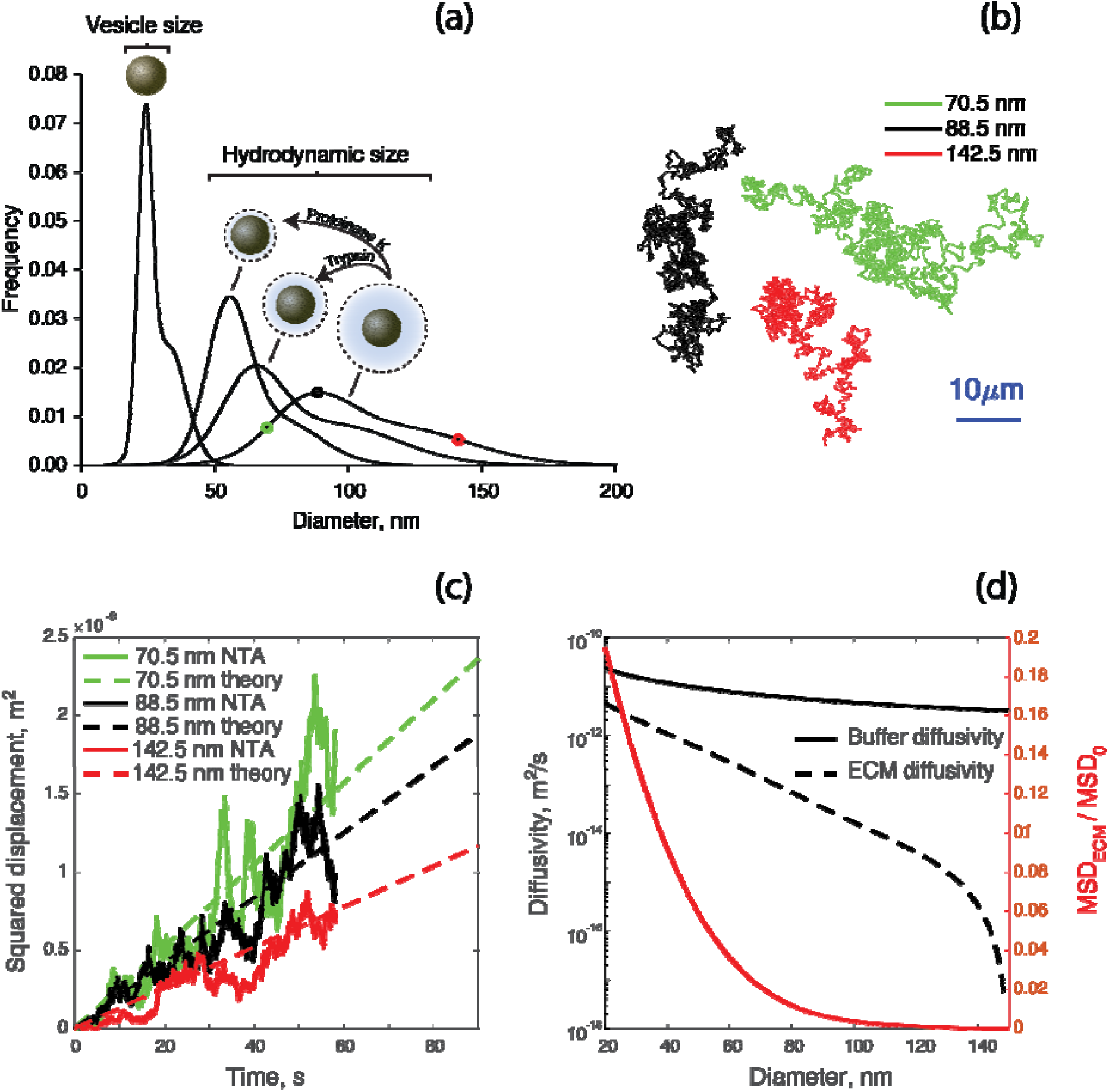
(a) Hydrodynamic sizes of exosomes are substantially larger than their vesicles. The proteolysis reduces the hydrodynamic size, implicating surface proteins in the retardation of the exosome mobility. Non-specific protein digestion by proteinase K shifts the hydrodynamic diameters into the range of vesicle sizes. (b) The migration of the exosomes having the mode, 10th, and 90th percentile hydrodynamic diameters (color coded in panel a) was measured for ~ 58 seconds in PBS by the single-particle tracking. (c) The squared particle displacements from the initial position corresponding to the tracks in (b) and the theoretical MSD predictions extended beyond the observation period to indicate the long-term trend in size-dependent reach of exosomes migrating by diffusion. (d) In the ECM, the size dependence of the exosomal migration is more pronounced, as indicated by a more rapid decrease in the diffusivity with the hydrodynamic diameter compared to the buffer diffusivity. The exosomes are immobilized as their hydrodynamic diameter approaches the size of the ECM pore.

## DISCUSSION

The enzyme-dependence in the change of the exosome mobility after the proteolysis implicates the surface proteins as the cause of the retarded diffusivity of exosomes relative to the expectation based on the size of their membrane envelope and points to the thinning of the protein corona as the mechanism responsible for the observed reduction in the hydrodynamic diameter of digested exosomes (Fig. 6a). The significant resistance imposed by the surface proteins on the motion of exosomes has several implications. The considerable heterogeneity in hydrodynamic diameters reported here implies that the exosomes migrate through the cell microenvironment with a widely varying speed. The diverse migration rate was revealed from the observations of the exosome trajectories in the solution, which me measured by single-particle tracking. Three such trajectories are shown in Fig. 6b for the exosomes with 88.5, 70.5, and 142.5 nm hydrodynamic diameters, which correspond to the mode, 10th, and 90th percentiles of the size distribution in Fig. 6a. Smaller exosomes diffuse more readily, as quantified in Fig. 6c by the time-dependent squared displacement from the initial location, and are more likely to participate in signaling after overcoming the transport barrier to reach the recipient cell.

The migration of a secreted exosome in vivo terminates by either re-internalization into the releasing cell, degradation,^2^ or uptake by a proximal or distant cell. Termination may occur after entering and emerging from circulating blood, lymph, cerebrospinal, or another biofluid. In the sequence from an exosome secretion to termination, the migration through the extracellular spaces (ECS) seems unavoidable.

Little is known about the diffusive migration of exosomes through the ECS. An early insight on the subject is derived from theoretical considerations, computer simulations, and experimental studies investigating the size and composition-dependent hindrance to the transit of nanoparticles and molecules through the ECS and the extracellular matrix (ECM) that occupies it. As summarized in several reviews,^47–50^ the size-dependent exclusion and filtering by ECM contribute to the preferential transit of small molecules and NPs.^47^ While the reduction in the diffusivity only slows the biodistribution of small particles, the larger particles may become trapped in the ECM.^51^ The mechanisms responsible for lower extracellular diffusivity include the steric hindrance from the constitutive network of polypeptides and polysaccharides; the electrostatic and other interactions with the ECM components; the localized increase in viscosity; and higher drag due to the wall effect. The same mechanisms are likely to influence the ECM migration of the exosomes in size and composition dependent manner.

Following Rusakov and Kullmann’s approach to modeling the ECM migration of molecules and NPs,^52^ we relate the effective diffusivity of the exosomes in the ECS, *D*_*e*_, and their unimpeded diffusivity in the solution, *D*_0_ as *D*_0_ =(*τ*_*g*_ *τ*_*v*_)^2^*D*_*e*_. The geometric tortuosity, *τ*_*g*_ describe the reduction in the migration rate due to a longer path an exosome must traverse through a meandering extracellular pore. Theoretical considerations indicate that *τ*_*g*_ ≅ 1.4 for a diffusive transport in a porous medium approximation of the ECS formed by isotropically oriented parenchyma. For a straight cylindrical pore *τ*_*g*_ = 1. In our simulations, we used *τ*_*g*_ = 1.67 which provided the best fit to the experimental data on the tortuosity in the cortex of rats.^53^ The contribution of steric hindrance to the exosome migration, their specific and nonspecific (e.g., electrostatic) interactions with ECM, and all other factors impeding the movement of the exosomes through the ECS are then described by the "viscous" tortuosity *τ*_*v*_ In the absence of exosome-specific data, *τ*_*v*_ was modeled using a steric partition coefficient and the enhanced friction ratio of pore-to-solution friction coefficients,^54^ both of which depend on the ratio of the hydrodynamic diameter of the exosomes to the size of the ECS pore through which the diffusion occurs.

The ECM diffusivity in a 150 nm ECS pore, selected to represent the mean confining ECS dimension inside a brain of a living rat^55^ and modeled following the described approximation, was compared with unimpeded diffusivity in the solution. The model predicts the exaggerated impact of sizes on the exosome mobility after secretion into the ECS (Fig. 6d). Average distance traveled in the ECS by an exosome of the smallest sizes (in the range of membrane vesicle, Fig. 6a) is ~45% of the distance translocated in the buffer, as indicated by the ratio of the mean squared displacements (MSD) during hindered and free diffusion. This distance drops to ~10% for 80 nm particles and becomes negligible as the size of the exosomes approaches the pore size. Slower migration of hydrodynamically larger exosomes increases their dwell time in the proximity of a parent cell, which favors their trapping in the ECS. Trapped EV may reduce the permeability of the matrix, creating a positive feedback for exosome immobilization in the ECM.

In a recent report, the exosomes were found to promote extracellular aggregation of gliotoxic amyloid-β (Aβ) peptides associated with Alzheimer’s disease pathology, while the reduction in the exosome secretion by neural cells improved cognition.^56^ It seems that only trapped (not transitory) exosomes can provide extracellular agglomeration sites and contribute to the extracellular Aβ accumulation. The evidence that the EVs are routinely sequestered in the ECM was provided by Huleihel et al.^57^ who found matrix-bound nanovesicles in urinary bladder matrix and other ECM scaffolds of laboratory-produced and commercially-procured samples. After the enzymatic digestion, which released the trapped vesicles from the ECM, the membrane vesicles of the trapped EVs were found to be between 40 nm in diameter and up, while the corresponding modes of their hydrodynamic diameters were between ~80 to 130 nm for different samples. Exosomal surface biomarkers were not detected in this study, likely due to PK digestion used to release the vesicles from the ECM before the analysis. Nevertheless, the microRNA cargo indicates that, at least, some the trapped EVs are cell-secreted while the membrane and hydrodynamic sizes, which overlap with the exosomal range, points at the exosome contribution to the described population of the matrix-bound vesicles.

The ECM mobility model used to produce results shown in Fig. 6d views the dimension of the extracellular pore as a hard size cut-off on the hydrodynamic diameter above which the exosomes are unable to transit through the ECM. This simplification ignores the flexing of "soft" macromolecular corona to overcome steric hindrance imposed by the ECS and the ECM mesh network. It may be, therefore, more appropriate to define the cut-off size for exosome trapping based on the size of the membrane vesicles rather than their hydrodynamic diameter. The reduction of the coronal layer in vivo by extracellular enzymes, such as metalloproteinases, also argues in favor of using the size of the membrane envelope in defining the absolute cut-off size above which exosomes are trapped in the ECM. Interestingly, the proteolytic activity may be bidirectional, with the host matrix shown to be degraded by oncosomes (tumor exosomes), thus promoting motility of cancer cells^58^ and the exosomes secreted by them.

Several additional reciprocating ECM-EV influences occurring in vivo are unaccounted by the model. The extracellular microenvironment influences the conformation of integral and peripheral membrane proteins,^59^ which affects the size of the corona and, therefore, the mobility of the exosomes in the ECM. For example, the ionic strength of the extracellular fluid is a factor with multifaceted effects on protein conformation,^60^ zeta potential,^61^ membrane fluidity,^62^ and the modulation of electrostatic interactions with the extracellular mesh.^63^ The extracellular pH affects the coronal layer by modifying the lateral extension and the configuration of surface protein moieties.^64^ At lower pH, the protein backbones experience higher amplitude of transient backbone fluctuations,^65^ which should aid in conforming the coronal layer to the dimensions of a sterically hindered region, just as flexible molecules were shown to translate through the ECM while the diffusion of rigid molecules of similar molecular weight was restricted.^50^ Furthermore, just as the components of extracellular milieu adsorb onto the surfaces of synthetic nanoparticles^66–69^ and modify their hydrodynamic size,^70,71^ similar impact should be expected as exosomes migrate through the ECM.

Dynamic influences of the ECM on the coronal layer of the exosomes, the evidence obtained for inorganic and polymeric nanoparticles in complex biological fluids, and the similar results observed with synthetic membrane nanoparticles^72^ strongly suggests that the exosomal corona undergoes time and microenvironment-dependent transformations as exosomes migrate and circulate in vivo. Conformational changes, specific and non-specific adsorptions, proteolytic degradation, and glycosylation of surface proteins are examples of such transformations.^73^ The influence of the microenvironment on the exosome mobility suggests that their hydrodynamic size should be viewed as a *dynamic property* that may change with time and spatial translocation. As a practical matter, the common practice of reporting only the hydrodynamic size distribution of the exosomes provides an insufficient characterization of their morphological properties even at static conditions. At a minimum, the characterization of, both, mobility and vesicular sizes is advisable.

Several notable features of solid tumors present a particularly interesting case to study the interplay between the properties of the extracellular matrix and the exosome migration. Cancer cells in vitro tend to secrete exosomes at an elevated rate.^74^ The relative abundance of oncosomes in serum and other biofluids of cancer patients^75–78^ indicates that this higher secretion is accompanied by an increased flux of the exosomes through the ECM away from the tumor cells. Among the tumor microenvironment features facilitating the transit, the acidity of most solid tumors may enhance the mobility by altering the confirmation and the flexibility of membrane proteins. The characteristic rigidity of the tumor tissue, caused by a denser network of collagen fibers in tumor ECM,^79^ should impede the passage of the exosomes. However, the tumor ECM is not only stiff but also swollen due to the abundance of hyaluronan.^80^ Such swelling may rescue the exosome mobility because an elevated interstitial pressure of solid tumors^81^ likely aids in the dissemination of exosomes by convective pressure-driven drainage of the ECS content into blood vessels. Unlike the case of diffusive migration, in which coronal layer impeded random motion, in the convective flow the macromolecular decoration of exosome membranes entrances their motility in the direction of the flow. Furthermore, in the convective flow, the exosomes slowed by a collagen mesh or another obstacle will experience a persistent drag force in the flow direction. Such force may be sufficient to mechanically flex the coronal layer until it complies with the available pore space and the exosome escapes the confinement. In this case, the cut-off size above which an exosome is trapped in the ECM is likely to be smaller than the hydrodynamic size and closer to the size of the membrane envelope. The same convective drag mechanism may be in play in dislodging oncosomes immobilized by adsorption and other non-specific interactions. Drugs used to reduce interstitial pressure^82^ will reduce the convective drainage of interstitial fluid into the circulation, which may impede the trafficking of the oncosomes away from the tumor site and disrupt exosome-mediated pro-tumorigenic and prometastatic signaling.

The exosome-mediated signaling, the mechanisms of cellular uptake of the exosomes, and the role of surface ligands in this process are subjects of intensive research.^1,15,83,84^ Some aspects of the cellular uptake relevant to the effect of the microenvironment on the exosome mobility are beginning to emerge. Low pH in tumor microenvironment has been shown to correlate with higher efficiency of cellular uptake.^74^ Though the precise mechanism of the pH-dependent enhancement of the uptake remains unknown, changes in the conformations of membrane proteins at higher acidity will affect the backbone flexibility and hydrodynamic size of the exosomes during their approach and diffusion within the vicinity of the target cell prior to docking and uptake.

The importance of macromolecular surface decoration extends beyond its influence on exosome mobility. It was previously shown that exosomes treated with proteinase K, despite their higher mobility, are less likely to be internalized by cells,^85^ indicating the importance of surface proteins in exosome docking to the target and the subsequent cellular internalization. The combination of a small hydrodynamic diameter for a faster migration and an even smaller membrane envelope, indicating the presence of a macromolecular corona formed by surface proteins, appears advantageous for exosome-mediated cellular communication. Further studies are needed to elucidate the tangled roles the surface proteins are playing in controlling the mobility and the uptake of the exosomes.

## METHODS

### Cell culture

The MCF7 breast cancer cell line (American Type Culture Collection (ATCC), Manassas, VA) was stored in liquid nitrogen after delivery and before use. To subculture cells, the MCF7 cell line was thawed and plated on 150 mm plates along with the complete growth medium composed of the base Eagle’s Minimum Essential Medium (ATCC), 0.01 mg/mL human recombinant insulin and 10% exosome-free fetal bovine serum (System Biosciences, Mountain View, CA). The cell culture aeration was by 95% air and 5% CO_2_ at 37°C. Once the cells settled down, the media was changed (approximately 24 hours after plating). The plate was then split at 1:10 ratio and ten plates were cultured, each containing 20 mL of media. Media from 9 of these plates (180 mL) were harvested and pooled. Media was then divided into 60 mL and 120 mL, further split into 30 mL/tube and centrifuged at 3,000×g for 15 minutes. The supernatant was then transferred from each tube to a new sterile 50 mL tube, and the exosome isolation was performed.

### Human sera samples

Sera samples of a female pancreatic cancer patient and a healthy female were obtained from ARUP Laboratories Inc. (Salt Lake City, UT) and de-identified according to IRB protocol.

### Exosome isolation

Cell line exosomes were isolated by precipitation (ExoQuick-TC, System Biosciences (SBI), Mountain View, CA) following the manufacturer’s protocol. The cell medium was centrifuged at 3,000×g for 15 minutes to remove cells and cell debris. The precipitating solution was added (1:5 volume ratio), well mixed with the sample, refrigerated overnight, and then centrifuged at 1,500×g for 30 minutes at room temperature. After centrifugation, the supernatant was discarded. The remaining exosome pellet was spun for another 5 minutes at 1,500×g to separate the residual ExoQuick TC solution, which was removed without disturbing the pellet. The pellet was then re-suspended in PBS buffer and divided into multiple aliquots used in the described experiments.

Human exosomes were isolated from 1 mL serum using ExoQuick kit (SBI) using a similar procedure.

### Exosome biomarkers

The protein concentration in MCF7 exosome samples was measured using NanoDrop ND-8000 Spectrophotometer (Thermo Scientific, Wilmington, DE) and was found to be ~3.5 mg/mL. 600 μL of the exosomes lysis buffer (SBI) was added to 150 μL of exosome sample, containing approximately ~500 μg of exosome proteins. After vortexing for 15 sec, 9.4 mL of exosome array binding buffer, supplied with the Exo-Check antibody PVDF membrane array (SBI), was added. The antibody array was wetted in 5 mL distilled water for 2 min at room temperature. After decanting the water from the membrane, 10 mL of the exosome-lysate-binding mixture was pipetted onto Exo-Check membrane and incubated for 12 h on a shaker at 4° C. The mixture was then discarded, 10 mL of wash buffer was added and gently agitated on the membrane surface for 5 min at room temperature. After removing the wash buffer, 10 mL of detection buffer was pipetted on the membrane surface and incubated for 2 h on a rocker at room temperature. The detection buffer was then discarded, the surface was washed three times with the wash buffer, and 2 mL of the developer solution (SuperSignal West Femto Chemiluminescent Substrate, Thermo Scientific, Rockford, IL) was added to cover the array membrane completely. After 2 min, the developer solution was discarded, and the array’ s response was read using the Bio-Rad ChemiDoc XRS Imager System (Hercules, CA). The obtained grayscale image was parsed into 12 areas, which included two positive controls (a bright signal indicates that the detection reagents are working properly), a blank spot (establishes background signal), a negative control to characterize cellular contamination (cis-Golgi matrix protein, GM130), and eight surface proteins known to express to various degree on the surfaces of different exosome types (ANXA5, TSG101, FLOT1, ICAM, CD63, CD81, ALIX, and EpCAM). A positive expression of 3 of more of these proteins is expected to confirm EV enrichment.^86^ The maximum grayscale intensity within each of the 12 areas, was expressed as a percent value within the range between the minimum, defined by the blank spot (0% expression), and the maximum, equal to the average of two positive controls (100% expression). Intensities below that of the blank spot were assigned 0% expression.

### miRNA isolation and extraction

The success of the exosome isolation was confirmed by quantifying the content of onco-microRNA miR-21, known to be highly expressed in MCF7 exosomes, in the purified sample. The total microRNA extraction was performed by using the Total Exosome RNA and Protein Isolation Kit (Invitrogen, Carlsbad, CA) following manufacturer’s protocol. The final elution volume of 40 μL was combined with 200 μL of 2X Denaturing Solution, lysed during vortexing, and then incubated for 5 min on ice. 400 μL of Acid-Phenol:Chloroform was added and vortexed for 60 s. Samples were centrifuged (5 min at 10,000× g) to separate the mixture into aqueous and organic phases, and the upper aqueous phase was transferred to a fresh tube. 133 μL of 100% EtOH was added, vortexed, and transferred onto a column. After centrifugation (30 s at 10,000×g), the flow-through, which contains the small RNA, was combined with 300μ L of 100% EtOH, transferred onto a new column, and spun down (30 s at 10,000×g). The flow-through was discarded, and the column went through two washing steps after which it was placed into a fresh tube. The RNA was eluted in 40 μL RNase-free water and stored at -80 C for downstream RNA profiling by digital droplet PCR.

### miR-21 quantification by digital droplet PCR

TaqMan miRNA Assay was used to quantify miR-21 with dPCR. Applied Biosystems (Foster City, CA) provided the miR-21 TaqMan assay amplification primers and probes. The actual sequences are proprietary (assay ID# 4427975). dPCR reactions were prepared into a 25 μ L final volume by using 12.5 μ L TaqMan Fast Advanced Master Mix, 1μ L TaqMan Advanced miRNA Assay (20X), 0.5 μ L RNase-free water, and 1μ L Drop Stabilizer (RainDance Technologies, Lexington, MA). Droplets were generated on the Raindrop Source chip (RainDance Technologies) and eluted into PCR tubes. Amplification was done on a thermocycler using the following settings: 95° C for 10 min, 50 cycles of 95° C for 15 s, 58° C for 15 s, 60° C for 45 s (slow ramp speed 0.5° C/second), and finally 98° C for 10 min. After PCR, samples were transferred to the RainDrop Sense chip (RainDance Technologies) for single fluorescent droplet detection. The data were analyzed by using the RainDrop Analyst Software v.2 (RainDance Technologies). A count of negative (miR-21 absent) versus positive droplets (miR-21 present) was recorded. The results show the expected large number of miR-21 copies in MCF7 exosomes (Fig. S1B) and not in exosomes from healthy women (Fig. S1C).

### Nanoparticle tracking analysis

Exosome concentration and size distribution were measured after thawing the sample (stored at -80°C before the experiments) and allowing thermal equilibration to room temperature. The sample was diluted 1:500 with 1x PBS (measured pH=7.4) to keep the number of particle in the field of view of the NTA instrument in the range between approximately 40 and 80 particles. The concentration after dilution (typically between 7×10^8^ to 12×10^8^ particles/mL) was measured and the corresponding concentration in the original sample obtained after adjusting for the dilution.

The NTA measurements were performed by illuminating the sample with a 40-mW violet laser (405 nm wavelength; Nanosight model LM10, Malvern, Salisbury, UK). Each cell line (Aliquot 1 or 2) and serum sample were injected into the test cell using a 1 mL sterile syringe. The light scattered by the exosomes during their Brownian motion in the solution was video recorded with a high-sensitivity sCMOS camera (OrcaFlash2.8, Hamamatsu Photonics, Hamamatsu City, Japan). For each sample, a set of five different 60-second videos was recorded at 25 frames per second. The sample in the field of view of the instrument was refreshed with the syringe-induced flow between each recording. The camera was set to have the shutter speed of 30 ms, slider gain equal to 500, and the camera level at 15. During the NTA video analysis, the maximum jump mode and minimum track length were set to Auto, blur size was set to 1-pass, and the detection threshold for video processing was set to 6.

The obtained videos of the Brownian motion were analyzed using NTA software (Nanosight version 3.0). During the analysis, a spherical shape of the exosomes was assumed. The buffer viscosity was set equal to the viscosity of water. Throughout the nanoparticle tracking, manual temperature measurements inside the test cell remained at 20°C with maximum fluctuations of 0.1 degrees. The viscosity of water at such conditions stays nearly constant and equal to 1 cP.

Each analyzed video consisted of 1499 frames which captured tracks of between 6,000 to 10,000 particles, depending on exosome concentration in the sample. For each sample, the measurements were repeated five times, and the results were averaged to obtain the exosome concentration, their size distribution, mode, mean, and standard deviation. The consistency of measurements was characterized by the standard error of the mean (SEofM).

After the analysis, the sample was aspirated from the test cell and stored at 4°C for antibody array, surface protein digestion, TEM, and AFM analysis, all of which were performed on the same day as the NTA. The described NTA size quantification was applied to Aliquots 1 and 2 before and after the samples were subjected to the enzymatic digestion of soluble and surface proteins.

### Brownian Trajectories by Single Particle Tracking

Five consecutive one-minute NTA videos (640×480 pixels in each frame that captures 110×80 µm area of the sample) were used to visualize the Brownian motion of diffusing MCF7 exosomes before enzymatic digestion. Out of the ensemble of the particles seen in each frame (AllTracks Nanosight data), the IDs of those corresponding to the mode of the size distribution, its 10th, and 90th percentile (88.5±1, 70.5±1, and 172.5± nm, respectively) were identified and the *,(x,y*) coordinate of their motion in consecutive frames extracted by a custom MATLAB function. Each particle of interest remained in the field of view between 0.2 and 4 seconds and an average ~0.7 secs. These tracking data were used to obtain longer Brownian trajectories (1453 video frames, corresponding to 58.12 seconds of motion at the Nanosight acquisition rate; Fig. 6b) by randomly concatenating the change in the position of the particles of the same size between consecutive video frames. The dimensions of the field-of-view were used to convert pixel coordinates into *μm*. For particles of different sizes, the obtained two-dimensional Brownian trajectories were used to find the squared displacements with time. The results were compared with the theoretical expectations for the mean squared displacement, < *r*^2^ >, vs. time (Fig. 6c) calculated as < *r*^2^ > = 4 *D*_0_*t*, where *r* is the distance from the initial (*t* = 0) to the current particle location, and *D*_0_is the particle diffusivity in the buffer.

### Dynamic Light Scattering

The isolated serum exosomes were diluted 1:100,000 in DI water and filtered through 0.2 μ m syringe filters (Corning, Tewksbury, MA). Prior to the measurements, 1 mL of the sample preparation was placed into a low volume disposable sizing cuvette for analysis and given 5 minutes to reach 25°C. The DLS measurements were performed on a Malvern Zetasizer Nano ZS (Worcestershire, UK) at 173° angle which measures particles in the 0.3–10,000 nm size range. Water viscosity at 25°C (0.8872 cP) and the refractive index of the solution equal to 1.33 were used to interpret the measurements. The refractive index for exosomes was set to 1.35. Samples were analyzed in 3 repeats, each consisting of 12 scattering measurements. The acquired data were processed using a General Purpose Model implemented in the Zetasizer software to obtain the size distribution, its mean, and the standard deviation.

### Atomic Force Microscopy (AFM)

Before imaging, the exosomes, known to have a negative zeta potential, were electrostatically immobilized on a surface according to the following procedure. A negatively charged surface of a freshly cleaved mica disk (10 mm diameter, Ted Pella Inc., Redding, CA) was modified to impart a positive surface charge by exposing it for 10 seconds to 100 μL of 10 mM NiCl _2_ solution (Sigma-Aldrich, St. Louis, MO) at room temperature. The solution was then removed, and the surface was washed three times with DI water and gently dried with nitrogen gas. The modified mica disk was placed in a petri dish and 100 μL of exosome solution, obtained by diluting a small volume of Aliquot 1 with DI water (1:25 dilution), was pipetted on the surface. The sealed petri dish was incubated for 18 hours at 4°C to allow for the exosome adsorption on a positively charged surface. After the incubation, the remaining solution was removed by aspiration, and the mica surface was washed four times with DI water, taking care to ensure that the surface remained hydrated throughout the process. Up to this point, the preparation steps needed to immobilize exosomes on the surface were identical for the samples to be imaged in the hydrated state or after desiccation but diverged from here.

The AFM imaging of the exosomes immobilized on a surface in their innate hydrated form or after desiccation was performed by Veeco Multimode Nanoscope V controller (Veeco Instruments Inc., Town of Oyster Bay, NY). To image the sample in the liquid, ~ 40 μL of 1× PBS was pipetted on mica surface and 5× 5 µm area was scanned in 512 lines with MLCT triangular cantilever (175 μm nominal length, 22 µm width, 0.07 N/m spring constant, and 20 nm tip radius; Bruker, Billerica, MA) held in Bruker’s MTFML probe holder. The scan rate in the tapping mode was set to 0.8 Hz for all samples, with the drive frequency maintained at ~7 kHz.

The desiccated sample was obtained by drying the hydrated sample in nitrogen gas after its AFM imaging in liquid was completed. The AFM imaging in the air was performed using TESP rectangular cantilever (125 μm nominal length, 40 μm width, 42 N/m spring constant and 8 nm tip radius; Bruker, Billerica, MA) held in MFMA tip holder. The 5×5 µm area of the sample (512 lines) was scanned in the tapping mode at 0.8 Hz scan and the drive frequency of ~260 kHz.

The acquired AFM data for hydrated and dried samples were analyzed with Gwyddion data visualization and analysis software (ver. 2.45, Czech Metrology Institute, Czech Republic). *Plane Level* function that subtracts the computed plane from the raw data was first applied to the raw AFM images. Image rows were then aligned by using the median method of *Align Rows* function which finds the corresponding height of each scan line and subtracts the corresponding quantities from each row. To remove the contribution of the local fault of the closed loop (a common scanning error), *Remove Scars* function was applied that automatically uses neighborhood lines to fill the gaps and eliminate the scars. The next step was to identify the particles in the image by using *Mark Grains by Threshold* function with the height threshold value chosen to be ~20%. *Distribution of Various Grain Characteristics* was then used to obtain the data on the radius of circles fit to the boundary of the particles in the XY plane, the height of each particle, and its volume. The particle volume was determined as the volume between the grain surface and the basis surface, where the basis surface is obtained by Laplacian interpolation of the data points surrounding each grain. The result was used to estimate the innate hydrated size of the exosomes in the solution. The previously reported globular shape of the exosomes in the solution was assumed. The globular size of each imaged exosome was estimated by calculating the diameter of a sphere with the same volume as enveloped by the corresponding exosome immobilized on the changed mica surface and distorted into a highly oblate shape by electrostatic forces. The obtained size estimate for the hydrated sample is given in Fig. 3.

Figures S2A and S2B give the representative tapping-mode height and phase images of the surface-bound exosomes after they were desiccated on the mica surface (compare with the corresponding images of the hydrated exosomes in Fig. 2). The elevation above the mica surface along the line crossing four different exosomes is given in Fig. S2C. After the desiccation, the exosomes are even more oblate than in the hydrated state (compare Fig. S2D with Fig. 2 and 3). Their height above the surface, characterized as a distribution in Fig. S2E, is reduced compared to the hydrated state, while the occupied surface area is expanded, Fig. S2F. Surprisingly, even after an addition shape distortion brought about by the desiccation, the volume enveloped by the exosome membrane remains essentially the same as the volume in the hydrated state, Fig. S2G.

### Cryo-Transmission Electron Microscopy (Cryo-TEM)

Approximately 3.5 μL of each analyzed sample was placed on a holey carbon-coated copper grid, and vitrified with the aid of a robotic accessory (FEI Vitrobot, Hillsboro, OR) by plunging it into liquid ethane maintained at the temperature of liquid nitrogen. Prior to sample application, grids were glow-discharged to render them hydrophilic. After vitrification, the samples were stored in liquid nitrogen. Before imaging, the stored samples were transferred to a cryoholder (Gatan 626, Pleasanton, CA), which continuously maintained their temperature at approximately -180° C. Cryo-TEM images were obtained at 200kV using an FEI Tecnai F20 transmission electron microscope (Hillsboro, OR) coupled to an FEI Eagle CCD camera. MCF7 exosomes were sized by a human operator using Image Processing Toolbox (MathWorks, Natick, MA). Sera exosomes were analyzed by a custom MATLAB routine, which converted the grayscale images into a binary form to define the boundary of the exosomes. The projection area occupied by each exosome was approximated by an ellipse, the perimeter of which provided the best least squares fit of the exosome boundary. The diameter of each analyzed exosome was set to the geometric mean of the lengths of the major and minor axis of the fitted ellipse.

### TEM imaging

EM imaging of Aliquot 1 was performed before and after its enzymatic treatment. Negatively stained specimens were prepared by a standard method. Small volumes (~ 3.5 μL) of exosome solutions were placed on Formvar-carbon-coated copper mesh grids made for transmission electron microscopy. Prior to use, grids were glow-discharged to render them hydrophilic. Each solution was allowed to adhere to the grid for approximately 1-2 minutes before blotting with filter paper. A small droplet (~3.5 μL) of 1% uranyl acetate solution was then placed on the grid. This solution was blotted away with filter paper after ~20 seconds, and the specimen was allowed to dry. Before application of stain, some samples were also treated with an intermediate washing step of 2-3 seconds in deionized water followed by blotting with filter paper. All the steps were performed at room temperature. The dried specimen was imaged in a transmission electron microscope (JEM1400Plus, JEOL USA, Peabody, MA) at 120 kV. Images were recorded on a Gatan Orius camera (Gatan, Inc., Pleasanton, CA, USA).

### Cleavage of surface proteins

The extracellular domain of exosomal proteins was cleaved by trypsin or proteinase K digestion using one of 4 different protocols summarized in Table 2. Trypsinization of was performed by adding 5 μL of 0.25% Trypsin-EDTA to 30 μL of the MCF7 exosome sample and incubating the mixture for 20 minutes at 37° C. After incubation, 5 μL of growth medium (EMEM) was added to stop trypsin activity. The variation of the above procedure (Trypsinization Protocol B) involved longer (1 hour) incubation at 37°C and did not include the addition of the growth media to the solution before the NTA characterization of the exosome sizes.

The alternative digestion protocol used proteinase K treatment during which 10 μL of proteinase K was added to 30 μL of MCF7 exosome samples. The mixture was incubated for 1 hour at 37 ° C after which the proteinase K activity was stopped by heating the sample to 65 ° C and maintaining it at the elevated temperature for an additional 10 minutes (Proteinase K Treatment Protocol C). The variation of this protocol (Proteinase K treatment protocol D) involves the size characterization by the NTA without proteinase K deactivation by the elevated temperature. All digestion experiments were performed in duplicates.

### Exosome diffusion in the ECM

The reduction in the effective diffusivity, *D*_*e*_, of the exosomes in the extracellular space relative to the unimpeded diffusivity in the solution,*D*_0_, was determined by the geometric (*τ*_*g*_) and viscous (*τ*_*v*_) tortuosities: *D*_*e*_ = (*τ*_*g*_ *τ*_*v*_)^-2^*D*_0_. The geometric tortuosity, which does not change with exosome sizes, was set to 1.67 – the value determined by fitting experimental data for murine cortex.^53^ The viscous tortuosity was calculated as *τ*_*v*_ (Φ_1_Φ_2_)^−1/2^, where Φ_1_ = (1 - λ)^2^ is a steric partitioning coefficient and the correction factor Φ_2_ was approximated by a frequently used relationship Φ_2_ = 1 - 2.1044λ + 2.089λ^3^ - 0.948 λ ^5,54^ Here, *λ* is the ratio of hydrodynamic exosome diameters to the size of the ECM pores, which we set equal to 150 nm to represent an average confining ECM dimension experimentally determined by Godin et al.^55^ for naive rat brain tissue. The exosomes larger than 150 nm are trapped in the ECM, Fig. 6d. The unimpeded diffusivity was calculated from the Stokes-Einstein equation, *D*_0_ = *k*_*B*_*T*/(*3πηd*) where*k*_*B*_ is the Boltzmann’s constant, η is the dynamic viscosity of PBS buffer and room temperature *T*, and *d* is the hydrodynamic diameter.

## ASSOCIATED CONTENT

### Supporting Information

Supplemental information file includes details on the quantification of antibody array and comparisons with prior reports; the expression of miR-21 microRNA in MCF7 exosomes and a negative control (Fig. S1); the summary of the AFM analysis of the desiccated exosomes (Fig. S2); and the TEM images of the MCF-7 exosomes before and after the proteolysis. This file is available free of charge via the Internet at http://pubs.acs.org.

## AUTHOR INFORMATION

### Author Contributions

MS and VSC wrote the manuscript through contributions of all authors. All authors approved the final version of the manuscript.

## ABBREVIATIONS

Aβ: gliotoxic amyloid-β peptides
AFM: Atomic-force microscopy
ALIX: programmed cell death 6 interacting protein
ANXA5: Annexin A5 protein
CD63 and CD81: tetraspanin proteins encoded by the CD63 and CD81 genes respectively
DLS: dynamic light scattering
ECM: extracellular matrix
ECS: extracellular space
EpCAM: epithelial cell adhesion glycoprotein
ER+: estrogen-receptor-positive
EV: extracellular vesicles
FLOT1: Flotillin-1 protein
GM130: cis-Golgi matrix protein
ICAM: Intercellular Adhesion Molecule 1
MCF7: Michigan Cancer Foundation-7 breast cancer cell line
miR-21: a microRNA encoded by MIR21 gene
microRNA: short non-coding RNA molecule
MSD: mean squared displacements
NP: nanoparticle
NTA: nanoparticle tracking analysis
PBS: phosphate-buffered saline
PCR: polymerase chain reaction
pdf: probability density function
PK: proteinase K
SEofM: standard error of the mean
SI: Supporting Information
TEM: transmission electron microscopy
TSG101: tumor susceptibility gene 101 protein.

## TABLE OF CONTENTS GRAPHIC

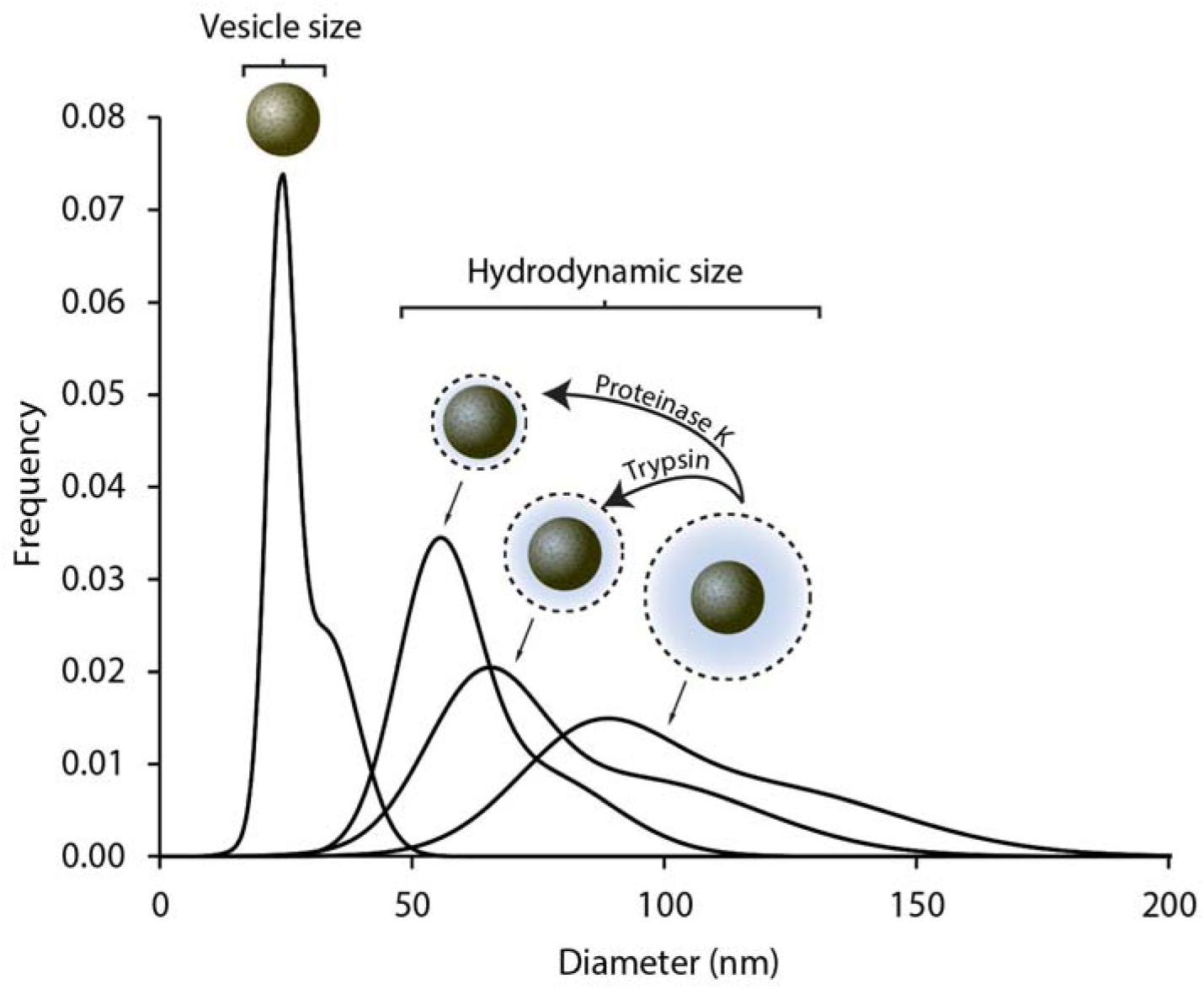

